# Evolutionary dynamics of collective action in spatially structured populations

**DOI:** 10.1101/012229

**Authors:** Jorge Peña, Georg Nöldeke, Laurent Lehmann

## Abstract

Many models proposed to study the evolution of collective action rely on a formalism that represents social interactions as *n*-player games between individuals adopting discrete actions such as cooperate and defect. Despite the importance of spatial structure in biological collective action, the analysis of *n*-player games games in spatially structured populations has so far proved elusive. We address this problem by considering mixed strategies and by integrating discrete-action *n*-player games into the direct fitness approach of social evolution theory. This allows to conveniently identify convergence stable strategies and to capture the effect of population structure by a single structure coefficient, namely, the pairwise (scaled) relatedness among interacting individuals. As an application, we use our mathematical framework to investigate collective action problems associated with the provision of three different kinds of collective goods, paradigmatic of a vast array of helping traits in nature: “public goods” (both providers and shirkers can use the good, e.g., alarm calls), “club goods” (only providers can use the good, e.g., participation in collective hunting), and “charity goods” (only shirkers can use the good, e.g., altruistic sacrifice). We show that relatedness promotes the evolution of collective action in different ways depending on the kind of collective good and its economies of scale. our findings highlight the importance of explicitly accounting for relatedness, the kind of collective good, and the economies of scale in theoretical and empirical studies of the evolution of collective action.

## 1 Introduction

Collective action occurs when individuals work together to provide a collective good (olson, 1971). Examples abound in the social and natural sciences: humans collectively build houses, roads, walls, and mobilize armies to make war; bacteria secrete enzymes that benefit other bacteria; sterile ant workers build the nest and raise the brood of the queen; lions work together to catch large game. Yet cooperation of this kind poses a collective action problem: if individual effort is costly there is an incentive to reduce or withdraw one’s effort, but if enough individuals follow this logic the collective good will not be provided. Much research in the social sciences has identified mechanisms for solving collective action problems, including privatization and property rights, reciprocity in repeated interactions, and institutions (Hardin, 1982; Sugden, 1986; Taylor, 1987; ostrom, 2003). The principles behind these mechanisms have also been explored in evolutionary biology (Boyd and Richerson, 1988; Nunn and Lewis, 2001; Strassmann and Queller, 2014) where it has been further emphasized that individual effort in cooperation should also increase as the relatedness between interactants increases (Hamilton, 1964). As social interactions often occur between relatives (because of spatial structure, kin recognition, or both; Rousset 2004; Bourke 2011) it is thought that relatedness plays a central role for solving collective action problems in biology. In particular, relatedness has been identified as the main mechanism of conflict resolution in the fraternal major transitions in evolution, i.e., those resulting from associations of relatives, such as the transitions from unicellularity to multicellularity, or from autarky to eusociality (Queller, 2000).

Mathematical models of collective action in spatially structured populations or between relatives often assume that strategies are defined in a continuous action space, such as effort invested into the provision of a public good or level of restrain in resource exploitation (Frank, 1995; Foster, 2004; Lehmann, 2008; Frank, 2010; Cornforth et al., 2012). This allows for a straightforward application of the direct fitness method (Taylor and Frank, 1996; Rousset, 2004) to investigate the effects of relatedness on the evolution of collective action. Contrastingly, many evolutionary models of collective action between unrelated individuals (Boyd and Richerson, 1988; Dugatkin, 1990; Motro, 1991; Bach et al., 2006; Hauert et al., 2006; Pacheco et al., 2009; Archetti and Scheuring, 2011; Sasaki and Uchida, 2014) represent interactions as *n*-player games in discrete action spaces (e.g., individuals play either “cooperate” or “defect”). These models can be mathematically involved, as identifying polymorphic equilibria might require solving polynomial equations of degree *n* − 1, for which there are no general analytical solutions if *n ≥* 6.

Here we integrate two-action *n*-player mixed strategy game-theoretic models into the direct fitness method of social evolution theory (Taylor and Frank, 1996; Rousset, 2004), which allows for studying the effect of spatial structure on convergence stability by using pairwise relatedness. Several shape-preserving properties of polynomials in Bernstein form (Farouki, 2012) then allow us to characterize convergence stable strategies with a minimum of mathematical effort. our framework delivers tractable formulas for games between relatives which differ from the corresponding formulas for games between unrelated individuals only in that “inclusive payoffs” (the payoff to self plus relatedness times the sum of payoffs to others) rather than solely standard payoffs must be taken into account. For a large class of games, convergence stable strategies can then be identified by a straightforward adaptation of existing results for games between unrelated individuals (Peña et al., 2014).

As an application of our modeling framework, we study the effects of relatedness on the evolution of collective action under different assumptions on the kind of collective good and its economies of scale, thus covering a wide array of biologically meaningful situations. To this aim, we distinguish between three kinds of collective goods: (i) “public goods” where all individuals in the group can use the good, e.g., alarm calls in vertebrates (Searcy and Nowicki, 2005) and the secretion of diffusible beneficial compounds in bacteria (Griffin et al., 2004; Gore et al., 2009; Cordero et al., 2012); (ii) “club goods” where only providers can use the good (Sandler and Tschirhart, 1997), e.g., cooperative hunting (Packer and Ruttan, 1988) where the benefits of a successful hunt go to individuals joining collective action but not to solitary individuals; and (iii) “charity goods” where only nonproviders can use the good, e.g., eusociality in Hymenoptera (Bourke and Franks, 1995) where sterile workers provide a good benefiting only queens.

For all three kinds of goods, we consider three classes of production functions giving the amount of good created as a function of the total level of effort and hence describing the associated economies of scale: (i) linear (constant returns to scale), (ii) decelerating (diminishing returns to scale), and (iii) accelerating (increasing returns to scale). Although linear production functions are often assumed because of mathematical simplicity, collective goods are often characterized by either decelerating or accelerating functions, so that the net effect of several individuals behaving socially is more or less than the sum of individual effects. In other words, social interactions can be characterized by (either positive or negative) synergy. For instance, enzyme production in microbial collective action is likely to be nonlinear, as in the cases of invertase hydrolyzing disaccharides into glucose in the budding yeast *Saccharomyces cerevisiae* (Gore et al., 2009) or virulence factors triggering gut inflammation in the *pathogen Salmonella typhimurium* (Ackermann et al., 2008). In the former case, the relationship between growth rate and glucose concentration in yeast has been reported to be decelerating, i.e., invertase production has diminishing returns to scale (Gore et al., 2009, Fig. 3.c); in the latter case, the relationship between the level of expression of virulence factors and inflammation intensity appears to be accelerating, i.e., it exhibits increasing returns to scale (Ackermann et al., 2008, Fig. 2.*d*).

We show that the effect of relatedness on the provision of collective goods, although always positive, critically depends on the kind of good (public, club, or charity) and on its economies of scale (linear, decelerating or accelerating production functions). Moreover, we show that relatedness and economies of scale can interact in nontrivial ways, leading to patterns of frequency dependence and dynamical portraits that cannot arise when considering any of these two factors in isolation. We discuss the predictions of our models, their implications for empirical and theoretical work, and their connections with the broader literature on the evolution of helping.

## 2 Model

### 2.1 Population structure

We consider a homogeneous group-structured population with a finite number of groups each containing an identical number of haploid individuals. Spatial structure may follow a variety of schemes, including the island model of dispersal (Wright, 1931), the isolation-by-distance model (Malécot, 1975), the haystack model (Maynard Smith, 1964), models where groups split into daughter groups and compete against each other (Gardner and West, 2006; Traulsen and Nowak, 2006; Lehmann et al., 2007b), and evolutionary graphs (ohtsuki et al., 2006; Taylor et al., 2007; Lehmann et al., 2007a). We leave particular details of the life history (e.g., whether generations are overlapping or non-overlapping) and population structure (e.g., the dispersal distribution) unspecified as they do not affect our analysis. All that is required is that the “selection gradient” can be written in a form proportional to (4) below. For this, we refer the interested reader to Rousset (2004); Lehmann and Rousset (2010); Van Cleve (2015).

### 2.2 Social interactions

Within groups, individuals participate in an *n*-player game with two available actions: *A* (e.g., “cooperation”) and *B* (e.g., “defection”). We denote by *a*_*k*_ the payoff to an *A*-player when *k* = 0, 1, …, *n* − 1 co-players choose *A* (and hence *n* − 1 − *k* co-players choose *B*). Likewise, we denote by *b*_*k*_ the payoff to a *B*-player when *k* co-players choose *A*. These payoffs can be represented as a table of the form:

| Opposing $A$-players | 0 | 1 | ... | $k$ | ... | $n-1$ |
| --- | --- | --- | --- | --- | --- | --- |
| payoff to $A$ | $a_0$ | $a_1$ | ... | $a_k$ | ... | $a_{n-1}$ |
| payoff to $B$ | $b_0$ | $b_1$ | ... | $b_k$ | ... | $b_{n-1}$ |

Individuals implement mixed strategies, i.e., they play *A* with probability *z* (and *B* with probability 1 − *z*). The set of available strategies is then the interval [0, 1]. At any given time only two strategies are present in the population: *z* and *z* + *δ*. Denoting by *z*_*•*_ the strategy of a focal individual and by *z*_*l*(*•*)_ the strategy of its *l*-th co-player, the expected payoff *π* to the focal can be written as

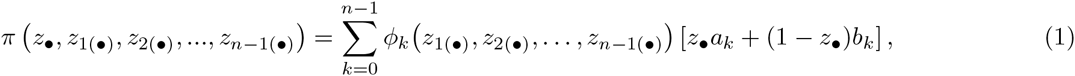

where *ϕ*_*k*_ is the probability that exactly *k* co-players play action *A*. A first-order Taylor-series expansion about the average strategy 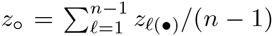 of co-players shows that, to first order in *δ*, the probability *ϕ*_*k*_ is given by a binomial distribution with parameters *n* − 1 and *z*_∘_, i.e.,

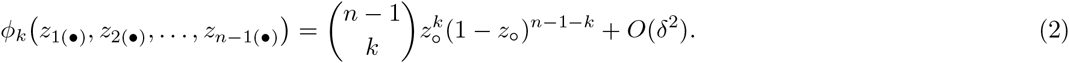

Substituting (2) into (1) and discarding second and higher order terms, we obtain

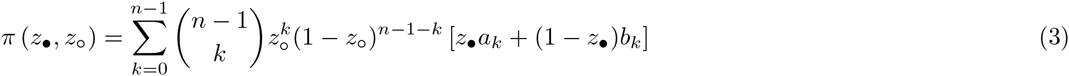

for the payoff of a focal individual as a function of the focal’s strategy *z*_*•*_ and the average strategy *z*_∘_ of co-players.

### 2.3 Evolutionary dynamics, scaled relatedness, and Hamilton’s rule

We are interested in the long-term evolutionary attractors of the probability *z* of playing *A*. To derive them, we consider a population of residents playing *z* in which a single mutant playing *z* + *δ* appears due to mutation, and denote by *ρ*(*δ, z*) the fixation probability of the mutant. We take the phenotypic selection gradient *S*(*z*) = (d*ρ/*d*δ*)_*δ*=0_ as measure of evolutionary success (Rousset and Billiard 2000, p. 819; Van Cleve 2015, Section 2.5); indeed, *S*(*z*) *>* 0 entails that the fixation probability of the mutant is greater than that of a neutral mutant under so-called “*δ*-weak” selection (Wild and Traulsen, 2007). Letting the expected relative fecundity of an adult be equal to its expected payoff (i.e., the payoffs from the game have fecundity effects; Taylor and Irwin 2000), the selection gradient *S*(*z*) can be shown to be proportional to what we call in this paper the “gain function”

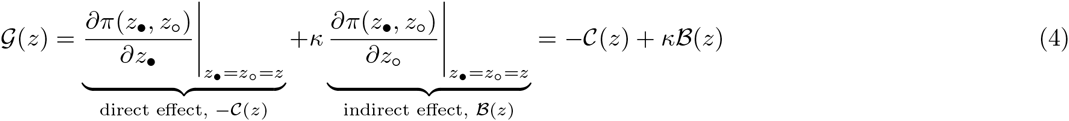

(for a derivation, see e.g., Van Cleve and Lehmann 2013, Eq. 7, or Van Cleve 2015, Eq. 73).

The gain function *𝒢*(*z*) is determined by three components. First, the direct effect −𝒞(*z*) describing the change in expected payoff resulting from the focal infinitesimally changing its own strategy. Second, the indirect effect *𝔅*(*z*) describing the change in expected payoff of the focal resulting from the focal’s co-players changing their strategy infinitesimally. Third, the indirect effect is weighted by the scaled relatedness coefficient *κ*, which is a measure of relatedness between the focal individual and its neighbors, demographically scaled so as to capture the effects of local competition on selection (Queller, 1994; Lehmann and Rousset, 2010).

Scaled relatedness *κ* is a function of demographic parameters such as the migration rate, group size, and vital rates of individuals or groups, but is independent of the evolving trait *z* and the payoffs from the game. In general, *κ* can take a value between −1 and 1, depending on the demographic assumptions (Lehmann and Rousset, 2010; Van Cleve and Lehmann, 2013). For instance, in a model where groups split into daughter groups and compete against each other (Traulsen and Nowak, 2006), scaled relatedness can be shown to be given by (Lehmann et al., 2007b)

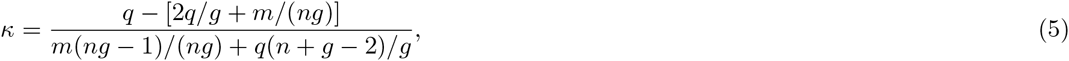

where *g* is the number of groups, *n* is group size, *q* is the splitting rate at which groups form propagules, and *m* is the migration rate (Van Cleve and Lehmann, 2013, Eq. B.4). Scaled relatedness coefficients have been evaluated for many spatially structured populations and demographic assumptions (see Lehmann and Rousset 2010; Van Cleve and Lehmann 2013 and references therein). In Appendix A we contribute to this literature by calculating values of scaled relatedness for several variants of the haystack model. In the subsequent analysis we treat *κ* as a parameter.

The gain function (4) is sufficient to characterize convergence stable strategies (i.e., strategies towards which selection locally drives the population by successive allelic replacements; Christiansen 1991; Geritz et al. 1998) under a trait substitution dynamic (Rousset and Billiard, 2000; Rousset, 2004). In our context, candidate convergence stable strategies are either singular strategies (i.e., values *z*^*\**^ for which *𝒢*(*z*^*\**^) = 0), or the two pure strategies *z* = 0 and *z* = 1. In particular, a singular strategy *z*^*\**^ is convergence stable (or an attractor) if d*𝒢*(*z*)*/*d*z|*_*z*=*z*\*_*<* 0 and convergence unstable (or a repeller) if d*𝒢*(*z*)*/*d*z|*_*z*=*z*\*_*>* 0. Regarding the endpoints, *z* = 0 (resp. *z* = 1) is convergence stable if *𝒢*(0) *<* 0 (resp. *𝒢*(1) *>* 0).

Finally, let us also note that the condition for a mutant to be favored by selection, −𝒞 + *κ𝔅 >* 0, can be understood as a demographically scaled form of the marginal version of Hamilton’s rule (Lehmann and Rousset, 2010), with *𝒞* corresponding to the marginal direct costs and *𝔅* to the marginal indirect benefits of expressing an increased probability of playing action *A*. This scaled version of Hamilton’s rule partitions the selection gradient in fecundity effects and scaled relatedness, in contrast to the partition on fitness effects and genetic relatedness of the classical formalism (i.e., −*c* + *rb >* 0, where *𝒞* and *b* are the direct and indirect fitness effects, and *r* is relatedness). Social evolution theory classifies social behaviors as altruistic, cooperative (or mutually beneficial), selfish, and spiteful, according to the signs of direct fitness costs and benefits (Hamilton, 1964; Rousset, 2004; West et al., 2007). A similar classification of social behaviors can be done according to the behavior’s effect on the direct and indirect components of marginal payoff (or fecundity). In order to avoid ambiguities, we refer to the resulting social behaviors as “payoff altruistic” (*𝒞 >* 0 and *𝔅 >* 0), “payoff cooperative” (*𝒞 <* 0 and *𝔅 >* 0), “payoff selfish” (*𝒞 <* 0 and *𝔅 <* 0), and “payoff spiteful” (*𝒞 >* 0 and *𝔅 <* 0).

## 3 Games between relatives

We start by deriving compact expressions for the direct effect *−𝒞*(*z*), the indirect effect *𝔅*(*z*), and the gain function *𝒢*(*z*) in terms of the payoffs *a*_*k*_ and *b*_*k*_ of the game. These expressions provide the foundation for our subsequent analysis.

Imagine a focal individual playing *B* in a group where *k* of its co-players play *A*. Suppose that the focal switches its action to *A* while co-players hold fixed their actions, thus changing its payoff from *b*_*k*_ to *a*_*k*_. As a consequence, the focal experiences a “direct gain from switching” given by

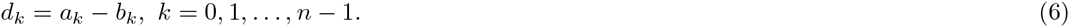

At the same time, each of the co-players playing *A* experiences a change in payoff given by Δ*a*_*k*−1_ = *a*_*k*_ − *a*_*k*−1_ and each of the co-players playing *B* experiences a change in payoff given by Δ*b*_*k*_ = *b*_*k*+1_ − *b*_*k*_.

Taken as a block, co-players experience a change in payoff given by

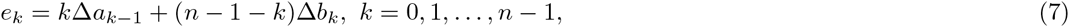

where we set *a*_−1_ = *b*_*n*+1_ = 0. From the focal’s perspective, this change in payoffs represents an “indirect gain from switching” to the focal if co-players are relatives. Adding up direct and indirect gains weighted by *κ* allows us to define the “inclusive gains from switching”

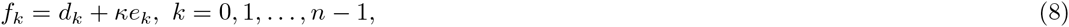

in a group where *k* out of the *n* − 1 co-players play *A*.

We show in Appendix B that the direct, indirect, and net effects appearing in (4) are indeed given by

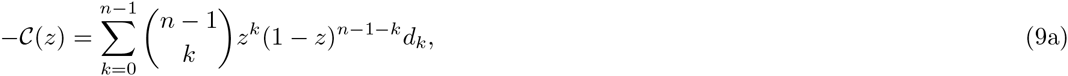

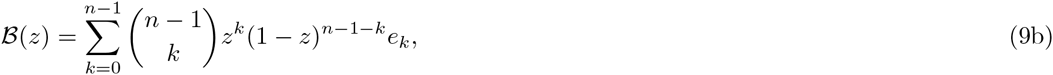

and

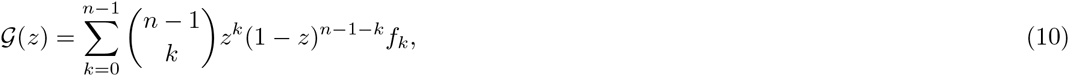

that is, as the expected values of the relevant gains from switching when the number of co-players playing *A* is distributed according to a binomial distribution with parameters *n* − 1 and *z*.

It follows from (10) that games between relatives are mathematically equivalent to transformed games between unrelated individuals, where “inclusive payoffs” take the place of standard, or personal, payoffs.

Indeed, consider a game in which *A*-players and *B*-players respectively obtain payoffs

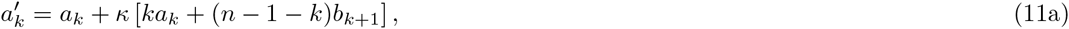

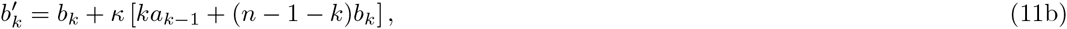

when *k* co-players play *A*. Payoffs 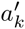 and 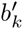 can be understood as inclusive payoffs consisting of the payoff obtained by a focal plus *κ* times the sum of the payoffs obtained by the focal’s co-players. Using (6)–(7) we can rewrite (8) as 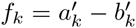, so that the inclusive gains from switching are identical to the direct gains from switching in a game with payoff structure given by (11).

This observation has two relevant consequences. First, existing results on the evolutionarily stable strategies of games between unrelated individuals (Peña et al., 2014), which are based on the observation that the right side of (10) is a polynomial in Bernstein form (Farouki, 2012), also apply here, provided that the inclusive gains from switching *f*_*k*_ are used instead of the standard (direct) gains from switching *d*_*k*_ in the formula for the gain function, and that evolutionary stability is understood as convergence stability. For a large class of games, these results allow us to identify convergence stable strategies from a direct inspection of the sign pattern of the inclusive gains from switching *f*_*k*_. Second, we can interpret the effect of relatedness as inducing the payoff transformation 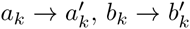. For *n* = 2, such transformation is the classic result of two-player games between relatives (Hamilton, 1971; Grafen, 1979; Day and Taylor, 1998)

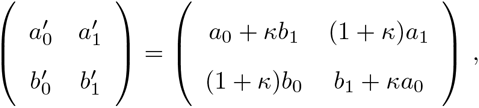

where the payoff of the focal is augmented by adding *κ* times the payoff of the co-player.

## 4 The evolution of collective action

Let us now apply our model to the evolution of collective action. To this end, we let action *A* (“provide”) be associated with some effort in collective action, action *B* (“shirk”) with no effort, and refer to *A*-players as “providers” and to *B*-players as “shirkers”. Each provider incurs a cost *γ >* 0 in order for a collective good of value *β*_*j*_ to be created, where *j* is the total number of providers. We assume that the collective good fails to be created if no individual works (*β*_0_ = 0), and that the value of the collective good *β*_*j*_ is increasing in the number of providers (Δ*β*_*j*_ = *β*_*j*+1_ − *β*_*j*_ *≥* 0). We distinguish between three kinds of collective goods, depending on which individuals have access to the good: (i) “public goods”, (ii) “club goods”, and (iii) “charity goods”. Fig. 1 illustrates these three kinds of collective goods and Table 1 provides the corresponding payoffs and gains from switching.

**Figure 1:**
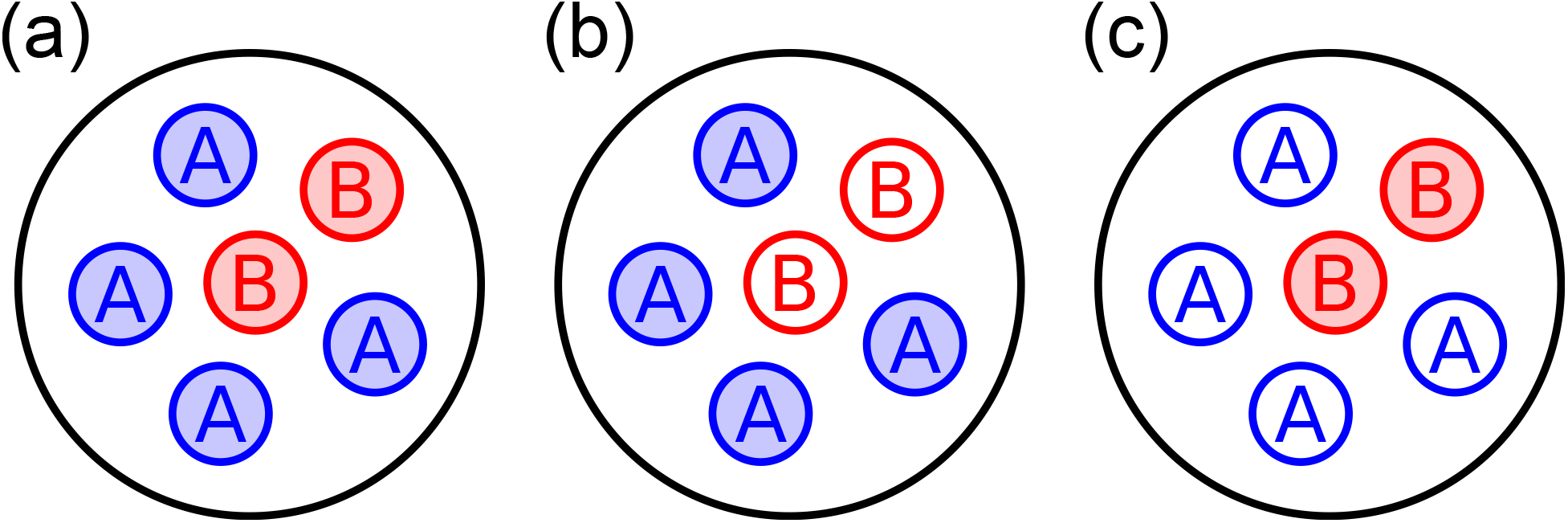
Three kinds of collective goods. Providers (*A*) and shirkers (*B*) interact socially. Providers (e.g., vigilants, cooperative hunters, or sterile workers) work together to create a collective good (e.g., alarm calls, increased hunting success, or nest defense), which can be used exclusively by a subset of individuals in the group (filled circles). Shirkers do not make any effort in collective action. *a*, Public goods (both providers and shirkers use the good). *b*, Club goods (only providers use the good). *c*, Charity goods (only shirkers use the good).

**Table 1:** Payoff structures and gains from switching for the three classes of collective action problems. In each case providers incur a cost *γ >* 0 to create a collective good of value *β*_*j*_*≥* 0, where *j* is the number of providers in the group. The number of providers experienced by a focal is *j* = *k* if the focal is a shirker (action *B*), and *j* = *k* + 1 if it is a provider (action *A*). Direct gains (*d*_*k*_) and indirect gains (*e*_*k*_) are calculated by substituting the formulas for *a*_*k*_ and *b*_*k*_ into (6) and (7). Inclusive gains from switching (*f*_*k*_) are then obtained from (8).

| kind of good | payoffs to $A$<br>($a_k$) | payoffs to $B$<br>($b_k$) | direct gains<br>($d_k$) | indirect gains<br>($e_k$) | inclusive gains<br>($f_k$) |
| --- | --- | --- | --- | --- | --- |
| public | $-\gamma + \beta_{k+1}$ | $\beta_k$ | $-\gamma + \Delta\beta_k$ | $(n-1)\Delta\beta_k$ | $-\gamma + (1 + \kappa(n-1))\Delta\beta_k$ |
| club | $-\gamma + \beta_{k+1}$ | 0 | $-\gamma + \beta_{k+1}$ | $k\Delta\beta_k$ | $-\gamma + \beta_{k+1} + \kappa k\Delta\beta_k$ |
| charity | $-\gamma$ | $\beta_k$ | $-\gamma - \beta_k$ | $(n-1-k)\Delta\beta_k$ | $-\gamma - \beta_k + \kappa(n-1-k)\Delta\beta_k$ |

Economies of scale are incorporated in the model through the properties of the production function *β*_*j*_. We investigate three functional forms (Fig. 2): (i) linear (*β*_*j*_ = *βj* for some *β >* 0, so that Δ*β*_*j*_ is constant), (ii) decelerating (Δ*β*_*j*_ is decreasing in *j*), and (iii) accelerating (Δ*β*_*j*_ is increasing in *j*). We also say that returns to scale are respectively (i) constant, (ii) diminishing, or (iii) increasing. To illustrate the effects of economies of scale, we consider the “geometric production function”:

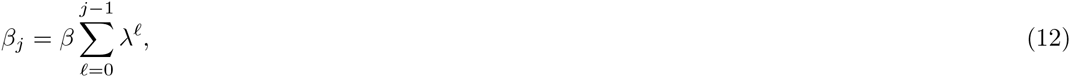

with *β >* 0 and *λ >* 0, for which returns to scale are constant when *λ* = 1, decreasing when *λ <* 1, and increasing when *λ >* 1 (Fig. 2).

**Figure 2:**
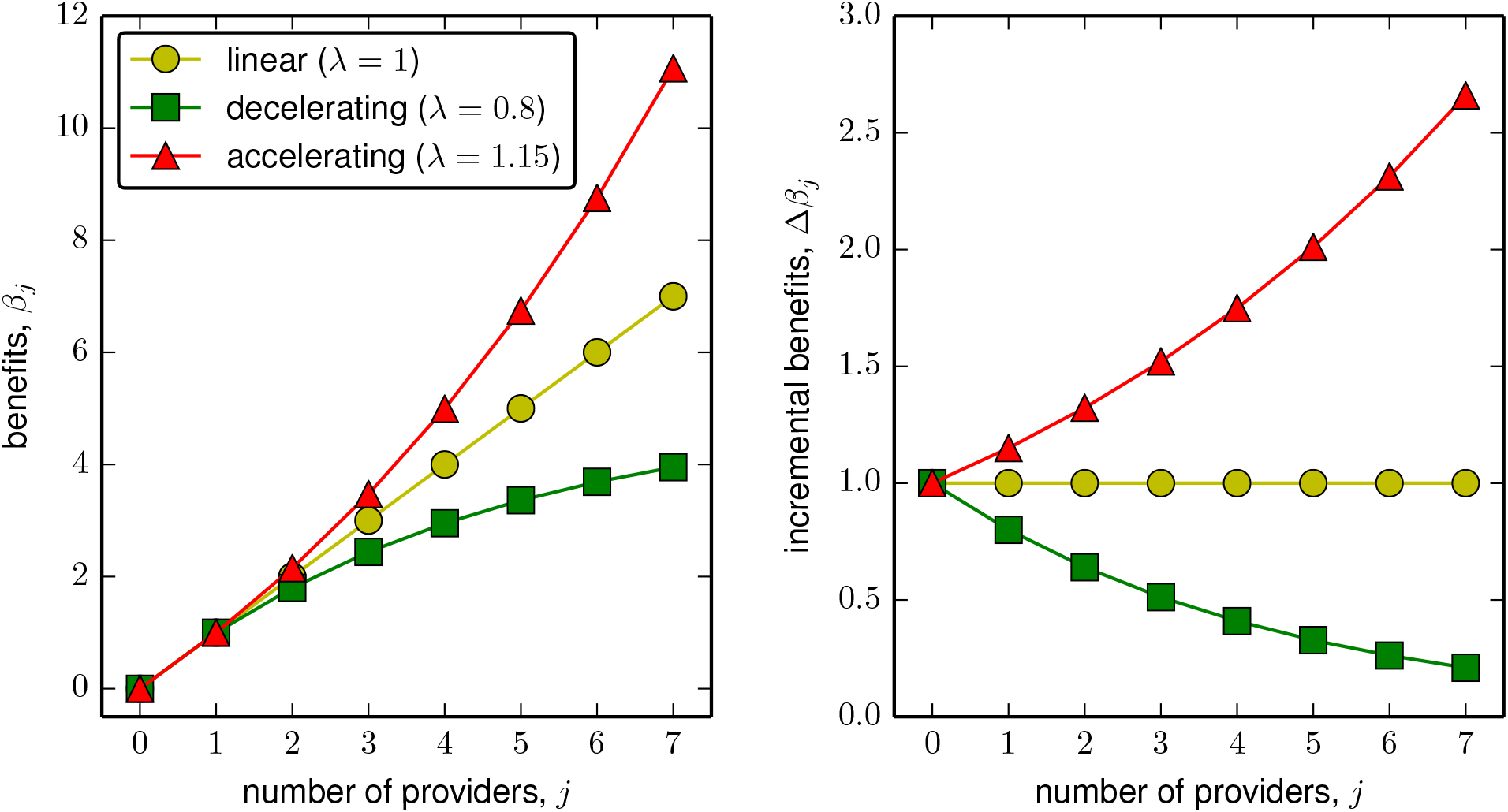
Linear, decelerating and accelerating production functions (here, geometric production functions as given by (12) with different values for the returns-to-scale parameter *λ*). *Left panel*, benefits *β*_*j*_ from the collective good are additive for linear functions, subadditive for decelerating functions and superadditive for accelerating functions. *Right panel*, incremental benefits Δ*β*_*j*_ from the collective good are constant for linear functions (constant returns to scale), decreasing for decelerating functions (diminishing returns to scale), and increasing for accelerating functions (increasing returns to scale).

For all three kinds of collective goods, the indirect gains from switching are always nonnegative, hence the indirect effect *𝔅*(*z*) is nonnegative for all *z*. Consequently, participation in collective action is either payoff altruistic or payoff cooperative, and the selection gradient is increasing in *κ*. The provision of each kind of collective good however leads to a different collective action problem, as it is reflected in the different payoff structures of the corresponding games (Table 1). In particular, while the provision of charity goods is payoff altruistic for all *z*, the provision of public and club goods can be either payoff altruistic or payoff cooperative, depending on the parameters of the game and the resident strategy *z*.

In the following, we characterize the evolutionary dynamics of each of these three kinds of collective action problems and investigate the effects of (scaled) relatedness on the set of evolutionary attractors. Although many of our results also extend to the case of negative relatedness, for simplicity we restrict attention to nonnegative relatedness (*κ ≥* 0). It will be shown that the evolutionary dynamics fall into one of the following five dynamical regimes: (i) “null provision” (*z* = 0 is the only attractor), (ii) “full provision” (*z* = 1 is the only attractor), (iii) “coexistence” (there is a unique singular strategy *z*^*\**^ which is attracting), (iv) “bistability” (*z* = 0 and *z* = 1 are both attracting, with a singular repeller *z*^*\**^ dividing their basins of attraction), and (v) “bistable coexistence” (*z* = 0 is attracting, *z* = 1 is repelling, and there are two singular strategies *z*_L_ and *z*_R_, satisfying *z*_L_ *< z*_R_, such that *z*_L_ is a repeller and *z*_R_ is an attractor). Regimes (i)-(iv) are those classical from 2 *×* 2 games (Cressman, 2003, Section 2.2), while bistable coexistence can only arise for interactions with more than two players (indeed, bistable coexistence requires the polynomial *𝒢*(*z*) to have two sign changes, which is only possible if *n >* 2; Broom et al. 1997; Gokhale and Traulsen 2014).

### 4.1 Linear production functions

To isolate the effects of the kind of collective good, we begin our analysis with the case where the production function takes the linear form *β*_*j*_ = *βj*, i.e., *λ* = 1 in (12). For all three kinds of collective goods, the gain function can then be written as

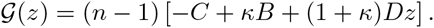

The parameter *C >* 0 may be thought of as the “effective cost” per co-player of joining collective action alone. We have *𝒞* = *γ/*(*n* − 1) when a focal provider is not among the beneficiaries of the collective good (charity goods) and *𝒞* = (*γ − β*)*/*(*n* − 1) otherwise (public and club goods). The parameter *B ≥* 0 measures the incremental benefit accruing to each co-player of a focal provider when none of the co-players joins collective action. We thus have *B* = 0 for club goods and *B* = *β* otherwise. Finally, *D* is null for public goods (*D* = 0), positive for club goods (*D* = *β*), and negative for charity goods (*D* = *-β*).

Depending on the values of these parameters, we obtain the following characterization of the resulting evolutionary dynamics:

1. For public goods (*D* = 0) selection is frequency independent. There is null provision if *-C* + *κB <* 0, and full provision if *−C* + *κB >* 0.
2. For club goods (*D >* 0) selection is positive frequency-dependent. There is null provision if *−C* + *κB* + (1 + *κ*)*D ≤* 0, and full provision if *−C* + *κB ≥* 0. If *−C* + *κB <* 0 *< −C* + *κB* + (1 + *κ*)*D*, there is bistability: both *z* = 0 and *z* = 1 are attractors and the singular strategy

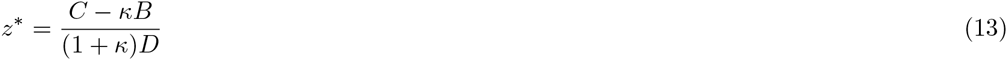

is a repeller.
3. For charity goods (*D <* 0), selection is negative frequency-dependent. There is null provision if *−C* + *κB ≤* 0, and full provision if *−C* + *κB* + (1 + *κ*)*D ≥* 0. If *−C* + *κB* + (1 + *κ*)*D <* 0 *< −C* + *κB*, there is coexistence: both *z* = 0 and *z* = 1 are repellers and the singular strategy *z*^*\**^ is the only attractor.

This analysis reveals three important points. First, in the absence of economies of scale the gain function is linear in *z*, which allows for a straightforward analysis of the evolutionary dynamics for all three kinds of collective action. Second, because of the linearity of the gain function, the evolutionary dynamics of such games fall into one of the four classical dynamical regimes arising from 2 *×* 2 games. Third, which of these dynamical regimes arises is determined by relatedness and the kind of good in a simple way. For all kinds of collective action, there is null provision when relatedness is low. For public goods provision, high values of relatedness lead to full provision. For club and charity goods, high relatedness also promotes collective action, leading to either bistability (club goods) or to the coexistence of providers and shirkers.

### 4.2 Public goods with accelerating and decelerating production functions

How do economies of scale change the evolutionary dynamics of public goods provision? Substituting the inclusive gains from switching given in Table 1 into (10) we obtain

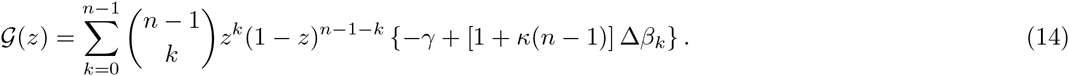

If the production function is decelerating, Δ*β*_*k*_ is decreasing in *k*, implying that *𝒢*(*z*) is decreasing in *z* (Peña et al., 2014, Remark 3). Similarly, if the production function is accelerating, Δ*β*_*k*_ is increasing in *k*, so that *𝒢*(*z*) is increasing in *z*. In both cases the evolutionary dynamics are easily characterized by applying existing results for public goods games between unrelated individuals (Peña et al., 2014, Section 4.3): with accelerating production functions, there is null provision if *γ ≥* [1 + *κ*(*n* − 1)]Δ*β*_0_, and full provision if *γ ≤* [1 + *κ*(*n* − 1)]Δ*β*_*n*−1_. If [1 + *κ*(*n* − 1)]Δ*β*_*n*−1_ *< γ <* [1 + *κ*(*n* − 1)]Δ*β*_0_, there is coexistence. With decelerating production functions, there is null provision if *γ ≥* [1 + *κ*(*n* − 1)]Δ*β*_*n*−1_, and full provision if *γ ≤* [1 + *κ*(*n* − 1)]Δ*β*_0_. If [1 + *κ*(*n* − 1)]Δ*β*_0_ *< γ <* [1 + *κ*(*n* − 1)]Δ*β*_*n*−1_, there is bistability.

The effect of relatedness on the evolution of public goods provision can be better grasped by noting that multiplying and dividing (14) by 1 + *κ*(*n* − 1) we obtain

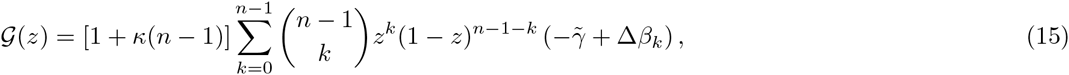

where 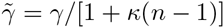. This is (up to multiplication by a positive constant) equivalent to the gain function of a public goods game with constant cost 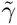 between unrelated individuals, which has been analyzed under different assumptions on the shape of the production function *β*_*k*_ (Motro, 1991; Bach et al., 2006; Hauert et al., 2006; Pacheco et al., 2009; Archetti and Scheuring, 2011; Peña et al., 2014). Hence, the effects of relatedness can be understood as affecting only the cost of cooperation, while leaving economies of scale and patterns of frequency dependence unchanged.

To illustrate the evolutionary dynamics of public goods games, consider a geometric production function (12) with λ ≠ 1 (see Table 2 for a summary of the results and Appendix C for a derivation). We find that there are two critical cost-to-benefit ratios:

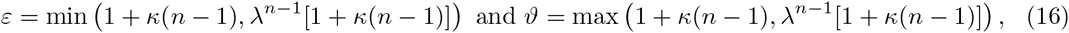

such that for small costs (*γ/β ≤ ε*) there is full provision and for large costs (*γ/β ≥ ϑ*) there is null provision. For intermediate costs (*ε < γ/β < ϑ*), there is a singular strategy given by

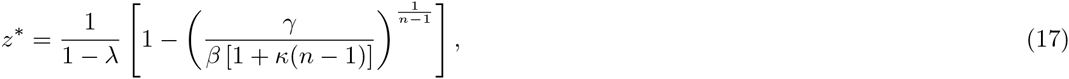

such that there is coexistence if returns to scale are diminishing (*λ <* 1) and bistability if returns to scale are increasing (*λ >* 1). For a given cost-to-benefit ratio *γ/β*, higher relatedness makes the region in the parameter space where cooperation (resp. defection) dominates larger (resp. smaller). Moreover, *z*^*\**^ is an increasing (resp. decreasing) function of *κ* when *λ <* 1 (resp. *λ >* 1), meaning that the proportion of providers at the internal attractor (resp. the size of the basin of attraction of *z* = 1) is larger for higher *κ* (Fig. 3.*a* and 3.*d*).

**Figure 3:**
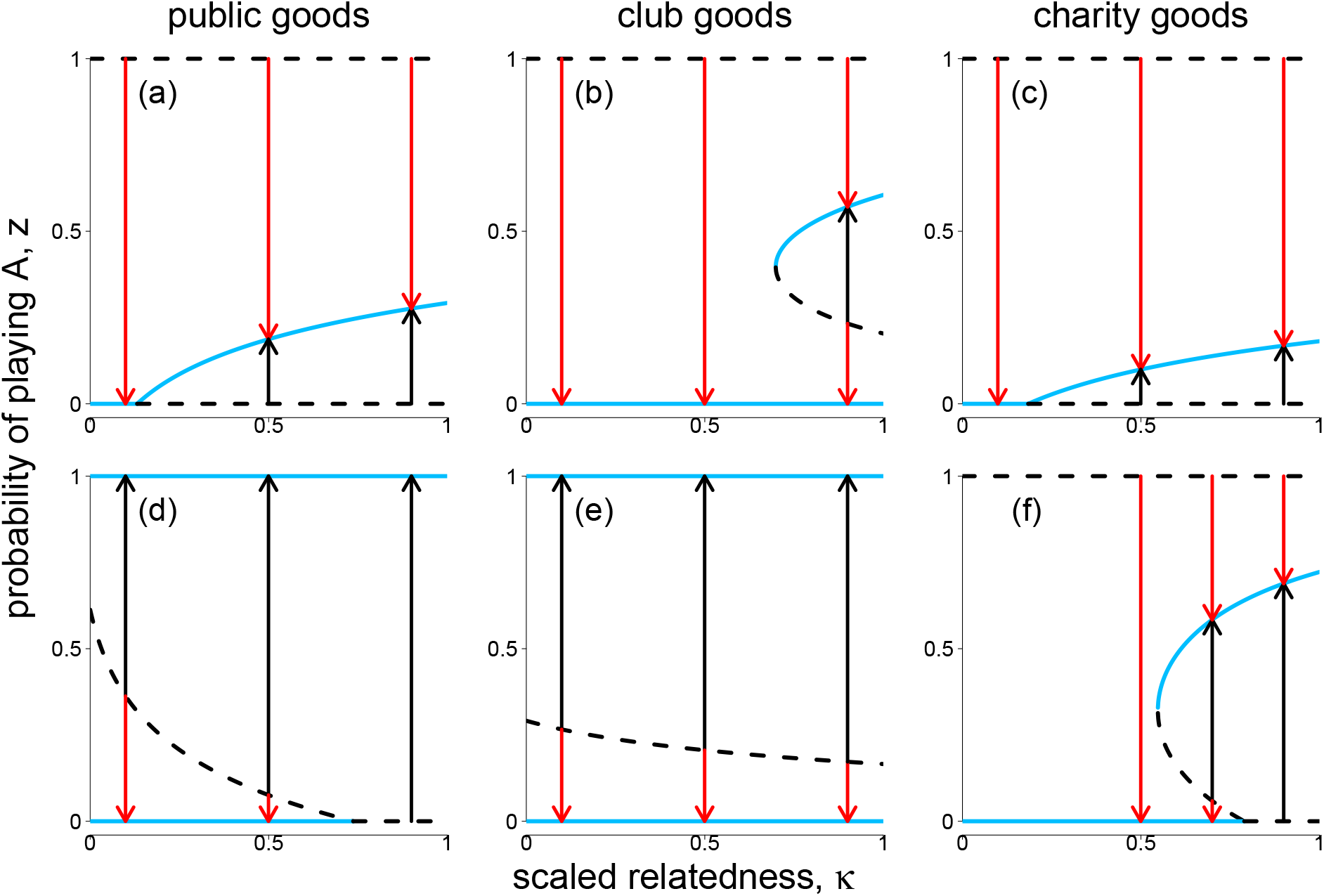
Bifurcation plots illustrating the evolutionary dynamics of collective action for public (*a*, *d*), club (*b*, *e*), and charity (*c*, *f*) goods with geometric production function. The scaled relatedness coefficient *κ ≥* 0 serves as a control parameter. Arrows indicate the direction of evolution for the probability of providing. Solid lines stand for convergence stable equilibria; dashed lines for convergence unstable equilibria. *a*, *b*, *c*, Diminishing returns to scale (*λ* = 0.7) and low cost-to-benefit ratio (*γ/β* = 3.5). *d*, *e*, *f*, Increasing returns to scale (*λ* = 1.25) and high cost-to-benefit ratio (*γ/β* = 15). In all plots, *n* = 20. The central arrows, for which *κ* = 0.5, could correspond, for example, to a group splitting model with infinitely many groups (*g ζ ∞*) and splitting probability equal to the migration rate *q* = *m* (5)*, or to a particular case of the haystack model with two founders (A.5)*.

**Table 2:** Dynamical regimes of collective action for the case of geometric production functions. The dynamical outcome depends on the type of good, the magnitude of the returns-to-scale parameter *λ*, and the cost-to-benefit ratio *γ/β*. The results hold for *κ ≥* 0 and *n >* 2. The critical cost-to-benefit ratios are given by *ζ* = *κ*(*n* − 1), *ε* = min (1 + *ζ, λ*^*n-*1^(1 + *ζ*)), *ϑ* = max(1 + *ζ, λ*^*n-*1^(1 + *ζ*)), *η* = [1*/*(*λ* − 1)] 1 + *λκ* [(*n* − 2)*λκ/*(1 + *ζ*)]^*n-*2^, *ς* = (1 − *λ*^*n*^)*/*(1 − *λ*) + *ζλ*^*n-*1^, *τ* = [1*/*(1 − *λ*)] 1 + *λκ* [(*n* − 2)*κ/*(1 + *ζ*)]^*n-*2^. The critical returns-to-scale parameters are *ξ* = *κ*(*n* − 2)*/*[1 + *κ*(*n* − 1)] and *ϱ* = 1*/ζ*.

| public goods | $\lambda < 1$ | | $\lambda > 1$ | |
| --- | --- | --- | --- | --- |
| | $\gamma/\beta \leq \varepsilon$ | full provision | $\gamma/\beta \leq \varepsilon$ | full provision |
| | $\varepsilon < \gamma/\beta < \vartheta$ | coexistence | $\varepsilon < \gamma/\beta < \vartheta$ | bistability |
| | $\gamma/\beta \geq \vartheta$ | null provision | $\gamma/\beta \geq \vartheta$ | null provision |
| club goods | $\lambda < 1/\varrho$ | | $\lambda \geq 1/\varrho$ | |
| | $\gamma/\beta \leq 1$ | full provision | $\gamma/\beta \leq 1$ | full provision |
| | $1 < \gamma/\beta < \varsigma$ | bistability | $1 < \gamma/\beta < \varsigma$ | bistability |
| | $\varsigma \leq \gamma/\beta < \tau$ | bistable coexistence | $\gamma/\beta \geq \varsigma$ | null provision |
| | $\gamma/\beta \geq \tau$ | null provision | | |
| charity goods | $\lambda \leq \varrho$ | | $\lambda > \varrho$ | |
| | $\gamma/\beta < \zeta$ | coexistence | $\gamma/\beta < \zeta$ | coexistence |
| | $\gamma/\beta \geq \zeta$ | null provision | $\zeta \leq \gamma/\beta < \eta$ | bistable coexistence |
| | | | $\gamma/\beta \geq \eta$ | null provision |

### 4.3 Club goods with accelerating and decelerating production functions

For club goods the direct gains from switching *d*_*k*_ (cf. Table 1) are increasing in *k* independently of any economies of scale. This implies that the direct effect −𝒞(*z*) is positive frequency-dependent. If the production function is accelerating, the indirect gains from switching *e*_*k*_ are also increasing in *k*, so that the indirect effect *𝔅*(*z*) is also positive frequency-dependent. For *κ ≥* 0 this ensures that, just as when economies of scale are absent, the gain function *𝒢*(*z*) is positive frequency-dependent. Hence, the evolutionary dynamics are qualitatively identical to those arising from linear production functions: for low relatedness, there is null provision; for high relatedness, there is bistability (see Fig. 3.*e* for an illustration and Appendix D.1 for proofs).

If the production function is decelerating, the indirect gains from switching *e*_*k*_ may still be increasing in *k* because the incremental benefit Δ*β*_*k*_ accrues to a larger number of recipients as *k* increases. In such a scenario, always applicable when *n* = 2, the evolutionary dynamics are again qualitatively identical to those arising when economies of scale are absent. A different picture emerges if the number of players is greater than two and returns to scale are diminishing. In this case, *𝔅*(*z*) can be negative frequency-dependent for some *z*, and hence (for sufficiently high values of *κ*) so can be *𝒢*(*z*). Depending on the value of relatedness, which modulates how the frequency dependence of *𝔅*(*z*) interacts with that of *𝒞*(*z*), and on the particular shape of the production function, this can give rise to evolutionary dynamics different from those discussed in Section 4.1. In particular, bistable coexistence is possible.

As an example, consider the geometric production function (12) with *λ /*= 1 (see Table 2 for a summary of results and Appendix D.2 for proofs). Defining the critical returns-to-scale value

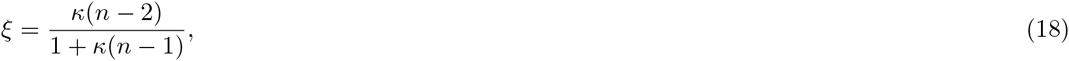

and the two critical cost-to-benefit ratios

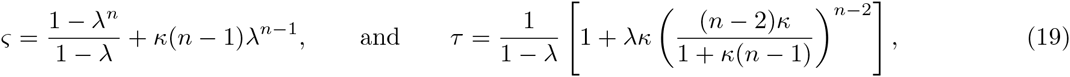

which satisfy *ξ <* 1 and *ς < τ*, our result can be stated as follows. For *λ ≥ ξ* the evolutionary dynamics depends on how the cost-to-benefit ratio *γ/β* compares to 1 and to *ς*. If *γ/β ≤* 1 (low costs), there is full provision, while if *γ/β ≥ ς* (high costs), there is null provision. If 1 *< γ/β < ς* (intermediate costs), there is bistability. For *λ < ξ*, the classification of possible evolutionary dynamics is as in the case *λ ≥ ξ*, except that, if *ς < γ/β < τ*, there is bistable coexistence, with *z* = 0 convergence stable, *z* = 1 convergence unstable, and two singular strategies *z*_L_ (convergence unstable) and *z*_R_ (convergence stable) satisfying 0 *< z*_L_ *< z*_R_ *<* 1. Although we have not been able to obtain closed form expressions for the singular strategies (*z*^*\**^ in the case of bistability; *z*_L_ and *z*_R_ in the case of bistable coexistence), numerical values of their locations can be obtained by searching for roots of *𝒢*(*z*) in the interval (0, 1), as we illustrate in Fig. 3.*b* and Fig. 3.*e*.

The critical values *ξ*, *ζ*, and *τ* are all increasing functions of *κ ≥* 0. Hence, with larger relatedness *κ*, the regions of the parameter space where some level of collective action is convergence stable expand at the expense of the region of dominant nonprovision. Moreover, inside these regions the convergence stable positive probability of providing increases with *κ* (Fig. 3.*b*). When the production function is “sufficiently” decelerating (*λ < ξ*) and for intermediate cost-to-benefit ratios (*ζ < γ/β < τ*), relatedness and economies of scale interact in a nontrivial way, leading to saddle-node bifurcations whereby two singular strategies appear as *κ* increases (Fig. 3.*b*).

### 4.4 Charity goods with accelerating and decelerating production functions

For charity goods the direct gains from switching *d*_*k*_ (cf. Table 1) are always decreasing in *k*, so that the direct effect −𝒞(*z*) is negative frequency-dependent.

From the formulas given in Table 1, it is clear that the direct gains from switching *d*_*k*_ are always decreasing in *k*. Hence, the direct effect −𝒞(*z*) is negative frequency-dependent. If the production function is decelerating, the indirect gains from switching *e*_*k*_ are also decreasing in *k*, implying that the indirect effect *𝔅*(*z*) is also negative frequency-dependent and that the same is true for the gain function *𝒢*(*z*) = −𝒞(*z*) + *κ𝔅*(*z*). Hence, diminishing returns to scale lead to evolutionary dynamics that are qualitatively identical to those arising when economies of scale are absent: for low relatedness, there is null provision, and for sufficiently high relatedness, a unique interior attractor appears (see Appendix E.1 and Fig. 3.*c*).

If the production function is accelerating, the indirect gains from switching *e*_*k*_ may still be decreasing in *k* because the incremental benefit Δ*β*_*k*_ accrues to a smaller number of recipients (*n* − 1 − *k*) as *k* increases. In such a scenario, always applicable when *n* = 2, the evolutionary dynamics are again qualitatively identical to those arising when economies of scale are absent. A different picture emerges if *n >* 2 holds and the economies of scale are sufficiently strong. In this case, *𝔅*(*z*) can be positive frequency-dependent for some *z*, and hence (for sufficiently high values of *κ*) so can be *𝒢*(*z*). Similarly to the case of club goods provision with diminishing returns to scale, this pattern of frequency dependence can give rise to bistable coexistence. For a concrete example, consider again the geometric production function (12) with *λ* 1 (see Table 2 for a summary of results, and Appendix E.2 for proofs). In this case, the evolutionary dynamics for *n >* 2 depend on the critical value

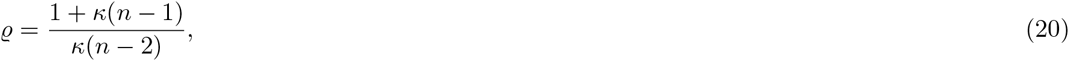

and on the two critical cost-to-benefit ratios

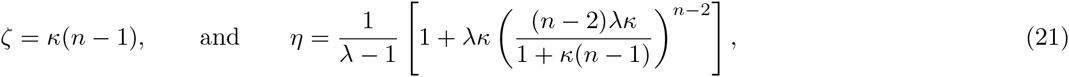

which satisfy *ϱ >* 1 and *ζ < η*.

With these definitions our results can be stated as follows. For *λ ≤ ϱ* the dynamical outcome depends on how the cost-to-benefit ratio *γ/β* compares to *ζ*. If *γ/β ≥ ζ* (high costs), there is null provision, while if *γ/β < ζ* (low costs), there is coexistence. For *λ > ϱ*, the dynamical outcome also depends on how the cost-to-benefit ratio *γ/β* compares to *η*. If *γ/β ≥ η* (high costs), there is null provision. If *γ/β ≤ ξ* (low costs), we have coexistence. In the remaining case (*ζ < γ/β < η*, intermediate costs) the dynamics are characterized by bistable coexistence. Closed form expressions for the singular strategies (*z*^*\**^ in the case of coexistence; *z*_L_ and *z*_R_ in the case of bistable coexistence) are not available, but we can find their values numerically, as we illustrate in Fig. 3.*c* and Fig. 3.*f*.

It is evident from the dependence of ϱ, *ζ*, and *η* on *κ* that relatedness plays an important role in determining the convergence stable level(s) of expression of helping. With higher *κ*, the regions of the parameter space where some *z >* 0 is convergence stable expand at the expense of the region of dominant nonprovision. This is so because *ζ* and *η* are increasing functions of *κ*, and ϱ is a decreasing function of *κ*. Moreover, inside these regions the stable non-zero probability of providing is bigger the higher *κ* (see Fig. 3.*c* and 3.*f*). Three cases can be more precisely distinguished as for the effects of increasing *κ*. First, *z* = 0 can remain stable irrespective of the value of relatedness, which characterizes high cost-to-benefit ratios. Second, the system can undergo a transcritical bifurcation, destabilizing *z* = 0 and leading to the appearance of a unique interior attractor (Fig. 3.*c*). This happens when *λ* and *γ/β* are relatively small. Third, there is a range of intermediate cost-to-benefit ratios such that, for sufficiently large values of *λ*, the system undergoes a saddle-node bifurcation, whereby two singular strategies appear (Fig. 3.*f*). In this latter case, economies of scale are strong enough to interact with the kind of good and relatedness in a nontrivial way.

### 4.5 Connections with previous models

Our formalization and analysis of specific collective action problems are connected to a number of results in the literature of cooperation and helping; we discuss these connections in the following paragraphs.

Our results on public goods games with geometric production functions (Section 4.2 and Appendix C) extend the model studied in (Hauert et al., 2006, p. 198) from the particular case of interactions between unrelated individuals (*κ* = 0) to the case of related individuals (*κ* ≠ 0) and recover the result in (Archetti, 2009, p. 476) in the limit *λ* → 0, where the game is known as the “volunteer’s dilemma” (Diekmann, 1985). Although we restricted our attention to the cases of linear, decelerating, and accelerating production functions, it is clear that (15) applies to production functions *β*_*j*_ of any shape. Hence, results about the stability of equilibria in public goods games with threshold and sigmoid production functions (Bach et al., 2006; Pacheco et al., 2009; Archetti and Scheuring, 2011; Peña et al., 2014) carry over to games in spatially structured populations.

Ackermann et al. (2008) considered a model of “self-destructive cooperation”, which can be reinterpreted as a charity goods game with no economies of scale in a particular version of the haystack model of population structure (Appendix A). In this model we have *κ* = (*n − N*)*/*(*n*(*N* − 1)), where *N* is the number of founders and *n ≥ N* is the number of offspring among which the game is played. Identifying our *γ* and *β* with (respectively) their *β* and *b*, the main result of Ackermann et al. (2008), given by Eq. 7 in their supplementary material, is recovered as a particular case of our result (13). The fact that in this example *κ* is a probability of coalescence within groups shows that social interactions effectively occur between family members, and hence that kin selection is crucial to the understanding of self-destructive cooperation (Gardner and Kümmerli 2008; see also Rodrigues and Gardner, 2013).

Eshel and Motro (1988) consider a model in which one individual in the group needs help, which can be provided (action *A*) or denied (action *B*) by its *n* − 1 neighbors: a situation Eshel and Motro call the “three brothers’ problem” when *n* = 3. Suppose that the cost for each helper is a constant *ε >* 0 independent on the number of volunteers (the “risk for each volunteer”, denoted by *𝒞* in Eshel and Motro 1988) and that the benefit for the individual in need when *k* co-players offer help is given by *v*_*k*_ (the “gain function”, denoted by *b*_*k*_ in Eshel and Motro 1988). Then, if individuals need help at random, the payoffs for helping (*A*) and not helping (*B*) are given by *a*_*k*_ = *−ε*(*n* − 1)*/n* + *v*_*k*_*/n* and *b*_*k*_ = *v*_*k*_*/n*. Defining *γ* = *ε*(*n* − 1)*/n* and *β*_*k*_ = *v*_*k*_*/*(*n* − 1), we have *a*_*k*_ = *−γ* + *β*_*k*_ and *b*_*k*_ = *β*_*k*_. Comparing these with the payoffs for public goods games in Table 1, it is apparent that the key difference between the case considered by Eshel and Motro (1988) and the public goods games considered here is that a provider cannot benefit from its own helping behavior. As we show in Appendix F, our results for public goods games carry over to such “other-only” goods games (Pepper, 2000). In particular, our results for public goods games with geometric benefits can be used to recover Results 1, 2, and 3 of Eshel and Motro (1988).

Finally, Van Cleve and Lehmann (2013) discuss an *n*-player coordination game. They assume payoffs given by *a*_*k*_ = 1 + *S*(*R/S*)^*k/*(*n*−1)^ and *b*_*k*_ = 1 + *P* (*T /P*)^*k/*(*n*−1)^, for positive *R, S, T*, and *P*, satisfying *R > T*, *P > S* and *P > T*. It is easy to check that both the direct effect −𝒞(*z*) and the indirect effect *𝔅*(*z*) are strictly increasing functions of *z* having exactly one sign change. This implies that, for *κ ≥* 0, the evolutionary dynamics are characterized by bistability. Importantly, and in contrast to the kinds of collective action analyzed in this article, expressing the payoff dominant action *A* does not always qualify as either payoff altruistic or payoff cooperative, as *𝔅*(*z*) is negative for some interval 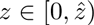. As a result, increasing relatedness *κ* can have mixed effects on the location of the interior convergence unstable equilibrium *z*^*\**^. Both of these predictions are well supported by the numerical results reported by Van Cleve and Lehmann (2013), where increasing *κ* leads to a steady increase in *z*^*\**^ for *R* = 2, *S* = 0.5, *P* = 1.5, *T* = 0.25, and a steady decrease in *z*^*\**^ for *R* = 2, *S* = 0.5, *P* = 1.5, *T* = 1.25 (see their Fig. 5). This illustrates the fact that scaled relatedness (and hence spatial structure) plays an important role not only in the specific context of collective action problems but also in the more general context of nonlinear *n*-player games.

## 5 Discussion

Many discrete-action, nonlinear *n*-player games have been proposed to study the evolutionary dynamics of collective action in well-mixed populations (Boyd and Richerson, 1988; Dugatkin, 1990; Motro, 1991; Bach et al., 2006; Hauert et al., 2006; Pacheco et al., 2009; Archetti and Scheuring, 2011; Peña et al., 2014). We extended these models to the more general case of spatially structured populations by integrating them into the direct fitness approach of kin selection theory (Taylor and Frank, 1996; Rousset, 2004; Lehmann and Rousset, 2010; Van Cleve, 2015). We showed that convergence stable strategies for games between relatives are equivalent to those of transformed games between unrelated individuals, where the payoffs of the transformed game can be interpreted as “inclusive payoffs” given by the original payoffs to self plus scaled relatedness times the sum of original payoffs to others. The evolutionary attractors of games in spatially and family structured populations can then be obtained from existing results on games in well-mixed populations (Peña et al., 2014).

We applied these general results to the evolution of collective action under different assumptions on the kind of collective good and its economies of scale, thereby unifying and extending previous analyses. We considered three kinds of collective goods, illustrative of different kinds of helping traits in nature. Firstly, public goods (both providers and shirkers have access to the good) for which the collective action problem is the well known free-rider problem (i.e., shirkers are cheaters who benefit from the good without helping to create it). Secondly, club goods (only providers have access to the good) for which there is no longer a free-rider but a coordination problem (i.e., individuals might prefer to stay alone rather than join a risky collective activity). Thirdly, charity goods (only shirkers use the good) for which the collective action problem takes the form of an altruism problem (i.e., individuals would prefer to enjoy the collective good rather than provide it for others).

We showed that relatedness can help solving each of these collective action problems, but that such effect takes different forms, depending on the kind of good and on its economies of scale. Simply put: relatedness transforms different collective action problems into different games. For public goods this transformation does not qualitatively affect the evolutionary dynamics, as it only reduces the cost of providing but otherwise leaves economies of scale (and hence patterns of frequency dependence) unaffected. Contrastingly, for club goods with diminishing returns and charity goods with increasing returns, relatedness can change patterns of frequency dependence in a nontrivial way. In particular, increasing relatedness can induce a saddle-node bifurcation resulting in the creation of an attracting equilibrium with positive helping and a repelling helping threshold.

This type of evolutionary dynamics, that we call bistable coexistence, is different from usual scenarios of frequency dependence in that selection favors mutants at some intermediate frequencies, but neither when rare nor common. Bistable coexistence had been previously predicted in models of public goods provision with sigmoidal production functions both in unstructured (Bach et al., 2006; Archetti and Scheuring, 2011) and structured (Cornforth et al., 2012) populations. our results show that bistable coexistence can also arise in models of club goods with diminishing returns and of charity goods with increasing returns when interactants are related. Participation in cooperative hunting illustrates the first of these situations: cooperative hunting is a club good (as hunted prey is available to hunters but not to solitary individuals) and is likely to exhibit diminishing returns because hunting success is subadditive in the number of hunters (Packer and Ruttan, 1988, Figs. 4-9). Eusociality in insects illustrates the second of these situations: eusociality is a charity good (as the benefits of the good created by workers are enjoyed only by reproducing queens) and is likely to exhibit increasing returns because of division of labor and other factors (Pamilo, 1991; Fromhage and Kokko, 2011). our results suggest that bistable coexistence might be more common than previously considered, thus expanding the repertoire of types of frequency-dependence selection beyond classic paradigms of either stabilizing (negative) or disruptive (positive) frequency-dependent selection (Levin et al., 1988).

Our results have implications for theoretical and empirical work on microbial cooperation. Although most research in this area has focused on public goods dilemmas (Griffin et al., 2004; Gore et al., 2009; Cordero et al., 2012), club and charity goods can also be present in microbial interactions. First, cases of “altruistic sacrifice” (West et al., 2006), “self-destructive cooperation” (Ackermann et al., 2008), and “bacterial charity work” (Lee et al., 2010), by which providers release chemical substances that benefit shirkers, are clear examples of charity goods. Second, “greenbeards” (Gardner and West, 2010; Queller, 2011), where providers produce an excludable good such as adherence or food sources (Smukalla et al., 2008; White and Winans, 2007), can be taken as examples of club goods. In all these examples, economies of scale are likely to be present, and hence also the scope for the complex interaction between relatedness and the shape of the production function predicted by our model. In particular, the possibility of bistable coexistence has to be acknowledged and taken into account both in models and experiments. Under bistable coexistence, even if providers are less fit than shirkers both when rare and when common, they are fitter than shirkers for some intermediate frequencies. Consequently, competition experiments should test for different starting frequencies before ruling out the possibility of polymorphic equilibria where providers and shirkers coexist. More generally, we encourage empirical work explicitly aimed at identifying club and charity goods and at measuring occurrences of economies of scale (i.e., nonlinear payoffs) in microbial systems.

We assumed that the actions implemented by players are discrete. This is in contrast to standard models of games between relatives, which assume a continuum of pure actions in the form of continuous amounts of effort devoted to some social activity. Such continuous-action models have the advantage that fitness or payoff functions (the counterparts to (3)) can be assumed to take simple forms that facilitate mathematical analysis. on the other hand, there are situations where individuals can express only few morphs (e.g., worker and queen in the eusocial Hymenoptera; Wheeler 1986), behavioral tactics (e.g., “producers” and “scroungers” in *Passer domesticus*; Barnard and Sibly 1981) or phenotypic states (e.g., capsulated and non-capsulated cells in *Pseudomonas fluorescens*; Beaumont et al. 2009). These situations are more conveniently handled by means of a discrete-action model like the one presented here. This being said, we expect our qualitative results about the interaction between kind of good, economies of scale, and relatedness to carry over to continuous-action models.

We assumed that the number of interacting individuals *n* is constant. However, changes in density will inevitably lead to fluctuating group sizes, with low densities resulting in small group sizes and high densities resulting in large group sizes. It is clear from the dependence of the critical cost-to-benefit ratios and the critical returns-to-scale parameters on group size (Table 2) that the effects of varying group sizes on the evolutionary dynamics of collective action will critically depend on the the kind of good and its economies of scale. It would be interesting to integrate this phenomenon into our model, thus extending previous work on the effects of group size in the evolution of helping (Motro, 1991; Brännström et al., 2011; Peña, 2012; Shen et al., 2014).

We assumed that players play mixed strategies and that the phenotypic deviation *δ* is small (i.e., “*δ*-weak” selection; Wild and Traulsen 2007), which is sufficient to characterize convergence stability but insufficient to characterize the fixation probability of a mutant when mutations have strong effects on phenotypes. This last scenario may occur when individuals can only express either full provision or null provision so that, say, mutants always play *A* and residents always play *B*. In these cases, a different limit of weak selection (i.e., “*w*-weak” selection; Wild and Traulsen 2007) might be more appropriate to model the evolutionary dynamics. For general nonlinear *n*-player games in structured populations the evolutionary dynamics will then depend not only on relatedness but also on higher-order genetic interactions (ohtsuki, 2014). The analysis of such evolutionary games under strong mutation effects and possibly strong selection remains to be done. This could be partly carried out by using invasion fitness proxies such as the basic reproductive number for subdivided populations (Metz and Gyllenberg, 2001; Ajar, 2003).

Collective action problems in nature are likely to be more diverse than the usually assumed model of public goods provision with constant returns to scale. Given the local demographic structure of biological populations, interactions between relatives are also likely to be the rule rather than the exception. Empirical work on the evolution of altruism and cooperation should thus aim at measuring the relatedness of interactants, the kind of good, and the associated economies of scale, as it is the interaction between these three factors which will determine the evolutionary dynamics of collective action in real biological systems.

## 6 Acknowledgements

This work was supported by Swiss NSF Grants PBLAP3-145860 (to JP) and PP00P3-123344 (to LL).

## A The haystack model

Many models of social evolution (Matessi and Jayakar, 1976; Wilson, 1987; Taylor and Wilson, 1988; Fletcher and Zwick, 2004; Ackermann et al., 2008; Powers et al., 2011; Cremer et al., 2012) have assumed variants of the haystack model (Maynard Smith, 1964), where several rounds of unregulated reproduction occur within groups before a round of complete dispersal. In these cases, as we will see below, the scaled relatedness coefficient *κ* takes the simpler interpretation of the coalescence probability of the gene lineage of two interacting individuals in their group. Here we calculate *κ* for different variants of the haystack model.

The haystack model can be seen as a special case of the island model where dispersal is complete and where dispersing progeny compete globally. In this context, the fecundity of an adult individual is the number of its offspring reaching the stage of global density-dependent competition. The conception of offspring may occur in a single or over multiple rounds of reproduction, so that a growth phase within patches is possible. We let *N* denote the number of founders (or lineages, or seeds) on a patch.

Two cases need to be distinguished when it comes to social interactions. First, the game can be played between the founders. In this case

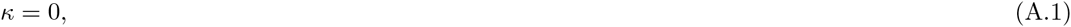

since relatedness is zero among founders on a patch and there is no local competition. Second, the game can be played between offspring after reproduction and right before their dispersal. In this case two individuals are related if they descend from the same founder. Since there is no local competition, *κ* is directly the relatedness between two interacting offspring and is obtained as the probability that the two ancestral lineages of two randomly sampled offspring coalesce in the same founder. (Relatedness in the island model is defined as the cumulative coalescence probability over several generations, see e.g., Rousset 2004, but owing to complete dispersal gene lineages can only coalesce in founders.)

In order to evaluate *κ* for the second case, we assume that, after growth, exactly *N*_o_ offspring are produced and that the game is played between them (*n* = *N*_o_). Founders, however, may contribute a variable number of offspring. Let us denote by *o*_*i*_ the random number of offspring descending from the founder *i* = 1, 2*, …, N* on a representative patch after reproduction, i.e., *O*_*i*_ is the size of lineage *i*. owing to our assumption that the total number of offspring is fixed, we have *N*_o_ = *O*_1_ + *O*_2_ + *…* + *O*_*N*_, where the *O*_*i*_’s are exchangeable random variables. The coalescence probability *κ* can then be computed as the expectation of the ratio of the total number of ways of sampling two offspring from the same founding parent to the total number of ways of sampling two offspring:

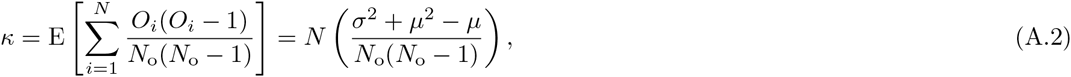

where the second equality follows from exchangeability, *µ* = E [*O*_*i*_] is the expected number of offspring descending from any founder *i*, and *σ*^2^ = E (*O*_*i*_ − *µ*)^2^ is the corresponding variance. Due to the fact that the total number of offspring is fixed, we also necessarily have *µ* = *N*_o_*/N* (i.e., *N*_o_ = E [*N*_o_] = E [*O*_1_ + *O*_2_ + *…* + *O*_*N*_] = *N µ*), whereby

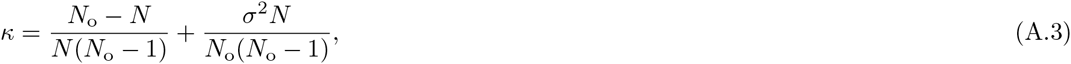

which holds for any neutral growth process.

We now consider three different cases:

1. Suppose that there is no variation in offspring production between founders, as in the life cycle described by Ackermann et al. (2008). Then *σ*^2^ = 0, and (A.3) simplifies to

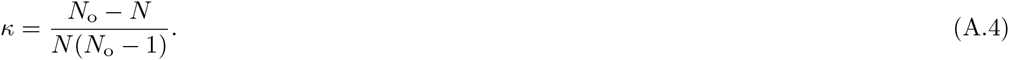
2. Suppose that each of the *N*_o_ offspring has an equal chance of descending from any founder, so that each offspring is the result of a sampling event (with replacement) from a parent among the *N* founders. Then, the offspring number distribution is binomial with parameters *N*_o_ and 1*/N*, whereby *σ*^2^ = (1 − 1*/N*)*N*_o_*/N*. Substituting into (A.3) we get

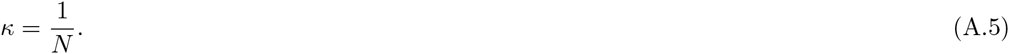 In more biological terms, this corresponds to a situation where individuals produce offspring according to a Poisson process and where exactly *N*_o_ individuals are kept for social interactions (i.e., the conditional branching process of population genetics; Ewens 2004).
3. Suppose that the offspring distribution follows a beta-binomial distribution, with number of trials *N*_o_ and shape parameters *α >* 0 and *β* = *α*(*N* − 1). Then, *µ* = *N*_o_*/N* and

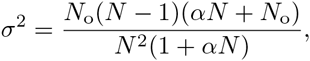

which yields

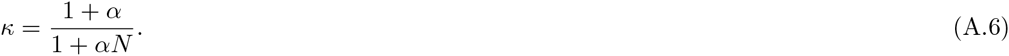

In more biological terms, this reproductive scheme results from a situation where individuals produce offspring according to a negative binomial distribution (larger variance than Poisson, which is recovered when *α → ∞*), and where exactly *N*_o_ individuals are kept for social interactions.

## B Gains from switching and the gain function

In the following we establish the expressions for −𝒞(*z*) and *𝔅*(*z*) given in (9); the expression for *𝒢*(*z*) (10) is then immediate from the definition of *f*_*k*_ (8) *and the identity G*(*z*) = −𝒞(*z*) + *κ𝔅*(*z*).

Recalling the definitions of *𝒞*(*z*) and *𝔅*(*z*) from (4) as well as the definitions of *d*_*k*_ and *e*_*k*_ from (6)–(7) we need to show

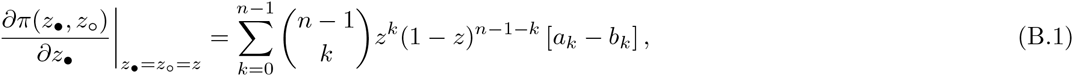

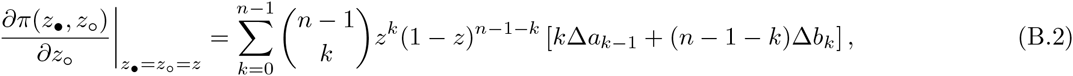

where the function *π* has been defined in (3). (B.1) follows directly by taking the partial derivative of *p* with respect to *z*_*•*_ and evaluating at *z*_*•*_ = *z*_∘_ = *z*, so it remains to establish (B.2).

Our derivation of (B.2) uses properties of polynomials in Bernstein form. Such polynomials, which in general can be written as 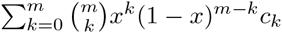 for *x ∈* [0, 1], satisfy (Farouki, 2012, p. 391)

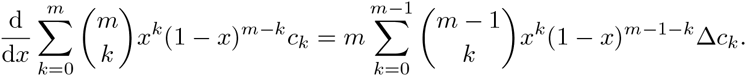

Applying this property to (3) and evaluating the resulting partial derivative at *z*_*•*_ = *z*_∘_ = *z*, yields

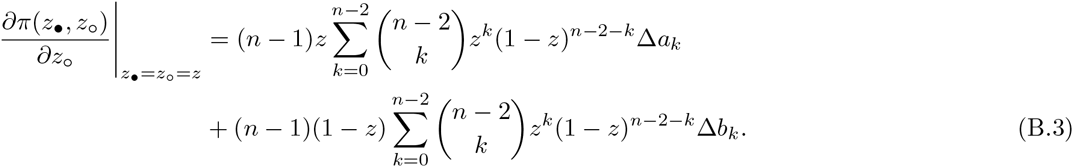

In order to obtain (B.2) from (B.3) it then suffices to establish

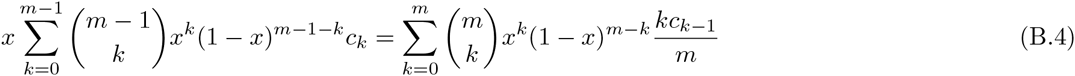

and

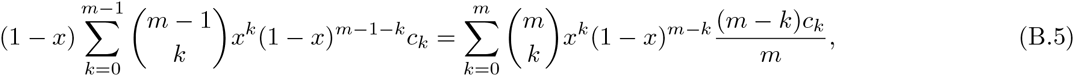

as applying these identities to the terms on the right side of (B.3) yields the right side of (B.2).

Let us prove (B.4) ((B.5) is proven in a similar way). Starting from the left side of (B.4), we multiply and divide by *m/*(*k* + 1) and distribute *x* to obtain

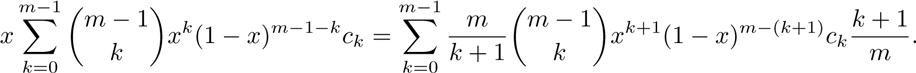

Applying the identity 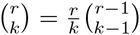 and changing the index of summation to *k* = *k* + 1, we get

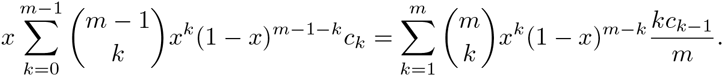

Finally, changing the lower index of the sum by noting that the summand is zero when *k* = 0 gives (B.4).

## C Public goods games with geometric production function

For a geometric production function, we have Δ*β*_*k*_ = *βλ*^*k*^, so that the inclusive gains from switching for public goods games are given by *f*_*k*_ = *−γ* + [1 + *κ*(*n* − 1)] *βλ*^*k*^. Substituting this expression into (10) and using the formula for the probability generating function of a binomial random variable, we obtain

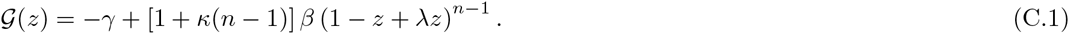

As *𝒢*(*z*) is either decreasing (*λ <* 1) or increasing (*λ >* 1) in *z*, *A* (resp. *B*) is a dominant strategy if and only if min [*𝒢*(0)*, 𝒢*(1)] *≥* 0 (resp. if and only if max [*𝒢*(0)*, 𝒢*(1)] *≤* 0). Using (C.1) to calculate *𝒢*(0) and *𝒢*(1) then yields the critical cost-to-benefit ratios *ε* = min [*𝒢*(0)*, 𝒢*(1)] and *ϑ* = max [*𝒢*(0)*, 𝒢*(1)] given in (16). The value of *z*^*\**^ given in (17) is obtained by solving *𝒢*(*z*^*\**^) = 0.

## D Club goods games

For club goods games, the inclusive gains from switching are given by

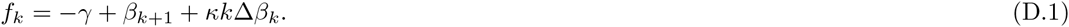

### D.1 Accelerating production function

In the case where the production function is accelerating, we have the following general result.

**Result 1** (Club goods games with accelerating production function). *Let f*_*k*_ *be given by* (D.1) *with β*_*k*_ *and* Δ*β*_*k*_ *increasing in k. Moreover, let κ ≥* 0*. Then*

1. *If γ ≤ β*_1_*, z* = 1 *is the only convergence stable strategy (full provision).*
2. *If β*_1_ *< γ < β*_*n*_ + *κ*(*n* − 1)Δ*β*_*n*−1_*, both z* = 0 *and z* = 1 *are convergence stable and there is a unique convergence unstable strategy z*^*\**^ *∈* (0, 1) *(bistability).*
3. *If γ ≥ β*_*n*_ + *κ*(*n* − 1)Δ*β*_*n*−1_*, z* = 0 *is the only convergence stable strategy (null provision).*

The assumptions in the statement of the result imply that *f*_*k*_ is increasing in *k*. In particular, we have *f*_0_ *< f*_*n*−1_. The sign pattern of the inclusive gain sequence thus depends on the values of its endpoints in the following way. If *f*_0_ *≥* 0 (which holds if and only if *γ ≤ β*_1_), *f*_*k*_ has no sign changes and a positive initial sign. If *f*_*n*−1_ *≤* 0 (which holds if and only if *γ ≥ β*_*n*_ + *κ*(*n* − 1)Δ*β*_*n*−1_), *f*_*k*_ has no sign changes and a negative initial sign. If *f*_0_ *<* 0 *< f*_*n*−1_ (which holds if and only if *β*_1_ *< γ < β*_*n*_ + *κ*(*n* − 1)Δ*β*_*n*−1_) *f*_*k*_ has one sign change and a negative initial sign. Result 1 follows from these observations upon applying Peña et al. 2014, Result 3.

### D.2 Geometric production function

For a geometric production function, we obtain the following result.

**Result 2** (Club goods games with geometric production function). *Let f*_*k*_ *be given by* (D.1) *with β*_*k*_ *given by* (12)*. Also, let κ ≥* 0 *and n >* 2 *(the cases κ <* 0 *or n* = 2 *are trivial). Moreover, let ξ, ζ and τ be defined by* (18) *and* (19)*. Then*

1. *If λ ≥ ξ, 𝒢*(*z*) *is nondecreasing in z. Furthermore*
  a. *If γ/β ≤* 1*, z* = 1 *is the only convergence stable strategy (full provision).*
  b. *If* 1 *< γ/β < ζ, both z* = 0 *and z* = 1 *are convergence stable and there is a unique convergence unstable strategy z*^*\**^ *∈* (0, 1) *(bistability).*
  c. *If γ/β ≥ ζ, z* = 0 *is the only convergence stable strategy (null provision).*
2. *If λ < ξ, 𝒢*(*z*) *is unimodal in z, with mode given by 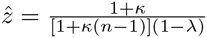. Furthermore*
  a. *If γ/β ≤* 1*, z* = 1 *is the only convergence stable strategy (full provision).*
  b. *If* 1 *< γ/β ≤ ζ, both z* = 0 *and z* = 1 *are convergence stable and there is a unique convergence unstable strategy 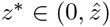(bistability).*
  c. *If ζ < γ/β < τ, there are two singular strategies z*_L_ *and z*_R_ *satisfying* 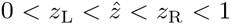. The strategies *z* = 0 *and z*_R_ *are convergence stable, whereas z*_L_ *and z* = 1 *are convergence unstable (bistable coexistence).*
  d. *If γ/β ≥ τ, z* = 0 *is the only convergence stable strategy (null provision).*

Observing that *ξ <* 1 holds and ignoring the trivial case *λ* = 1, there are three cases to consider: (i) *λ >* 1, (ii) 1 *> λ ≥ ξ*, and (iii) *ξ > λ*.

For *λ >* 1 the production function is accelerating and hence Result 1 applies with *β*_1_ = *β* and *β*_*n*_ + *κ*(*n* − 1)Δ*β*_*n*−1_ = *βζ*. This yields Result 2.1 for the case *λ >* 1.

To obtain the results for the remaining two cases, we calculate the first and second forward differences of the production function (12) and substitute them into

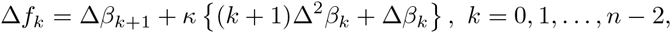

to obtain

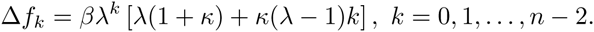

For *λ <* 1, the sequence Δ*f*_*k*_ is decreasing in *k* and hence can have at most one sign change. Moreover, as Δ*f*_0_ = *βλ*(1 + *κ*) *>* 0 always holds true, the initial sign of Δ*f*_*k*_ is positive and whether or not the sequence Δ*f*_*k*_ has a sign change depends solely on how Δ*f*_*n-*2_ compares to zero. observe, too, that for *λ <* 1 we have *ζ >* 1 as *λ*^*n*^ *< λ* holds.

Consider the case *ξ ≤ λ <* 1. By the definition of *ξ* (18) *this implies Δf*_*n-*2_ *≥* 0. In this case Δ*f*_*k*_ has no sign changes and *f*_*k*_ is nondecreasing. The sign pattern of the inclusive gain sequence can then be determined by looking at how the signs of its endpoints depend on the cost-to-benefit ratio *γ/β*. If *γ/β ≤* 1, then *f*_0_ *≥* 0, implying that *f*_*k*_ has no sign changes and its initial sign is positive. If *γ/β ≥ ζ*, then *f*_*n*−1_ *≤* 0 and hence *f*_*k*_ has no sign changes and its initial sign is negative. If 1 *< γ/β < ζ*, then *f*_0_ *<* 0 *< f*_*n*−1_, i.e., *f*_*k*_ has one sign change and its initial sign is negative. Result 2.1 then follows from an application of Peña et al. 2014, Result 3.

For *λ < ξ* we have Δ*f*_*n-*2_ *<* 0, implying that Δ*f*_*k*_ has one sign change from + to −, i.e., *f*_*k*_ is unimodal. Hence, the gain function *𝒢*(*z*) is also unimodal (Peña et al., 2014, Section 3.4.3) with mode 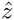 determined by 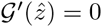. Using the assumption of geometric benefits, we can express *𝒢*(*z*) is closed form as

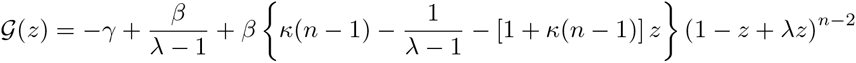

with corresponding derivative

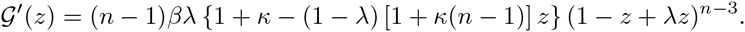

Solving 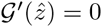 then yields 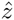 as given in Result 2.2. The corresponding maximal value of the gain function is

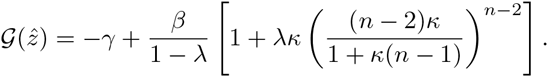

Result 2.2 then follows from applying Peña et al. 2014, Result 5. In particular, if *γ/β ≤* 1, we also have *γ/β < ζ*, ensuring that *f*_0_ *≥* 0 and *f*_*n*−1_ *>* 0 hold (with unimodality then implying that the gain function is positive throughout). If 1 *< γ/β ≤ ζ*, we have *f*_0_ *<* 0 and *f*_*n*−1_ *≥* 0 (with unimodality then implying 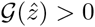. If *ζ < γ/β*, we have *f*_0_ *<* 0 and *f*_*n*−1_ *<* 0. Upon noticing that 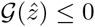 is satisfied if and only if *γ/β ≥ τ* holds, this yields the final two cases in Result 2.2.

## E Charity goods games

For charity goods games, the inclusive gains from switching are given by

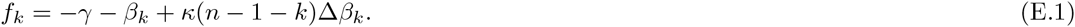

### E.1 Decelerating production function

If the production function is decelerating, we have the following general result.

**Result 3** (Charity goods games with decelerating production function). *Let f*_*k*_ *be given by* (E.1) *with β*_0_ = 0*, β*_*k*_ *increasing and* Δ*β*_*k*_ *decreasing in k. Moreover, let κ ≥* 0 *(the case κ <* 0 *is trivial). Then*

1. *If γ ≥ κ*(*n* − 1)Δ*β*_0_*, z* = 0 *is the only convergence stable strategy (null provision).*
2. *If γ < κ*(*n* − 1)Δ*β*_0_*, both z* = 0 *and z* = 1 *are convergence unstable and there is a unique convergence stable strategy z*^*\**^ *∈* (0, 1) *(coexistence).*

The arguments used for deriving this result are analogous to those used for deriving the results for the case of club goods with accelerating production function (Result 1 in Appendix D). The assumptions in the statement imply that *f*_*k*_ is decreasing in *k*. In particular, we have *f*_*n*−1_ *< f*_0_. Consequently, if *f*_0_ *≤* 0 (which holds if and only if *γ ≥ κ*(*n* − 1)Δ*β*_0_) the inclusive gain sequence has no sign changes and its initial sign is negative. observing that *f*_*n*−1_ = *−γ − β*_*n*−1_ *<* 0 always holds true, the inequality *f*_0_ *>* 0 (which holds if and only if *γ < κ*(*n* − 1)Δ*β*_0_) implies that the decreasing sequence *f*_*k*_ has one sign change and that its initial sign is positive. Result 3 is then obtained by an application of Peña et al. 2014, Result 3.

### E.2 Geometric production function

For a geometric production function, we obtain the following result.

**Result 4** (Charity goods games with geometric production function). *Let f*_*k*_ *be given by* (E.1) *with β*_*k*_ *given by* (12) *and let κ ≥* 0 *and n >* 2 *(the cases κ <* 0 *or n* = 2 *are trivial). Moreover, let ϱ, ζ and η be defined by* (20) *and* (21)*. Then*

1. *If λ ≤ ϱ, 𝒢*(*z*) *is nonincreasing in z. Furthermore:*
  a. *If γ/β < ζ, both z* = 0 *and z* = 1 *are convergence unstable and there is a unique convergence stable strategy z*^*\**^ *∈* (0, 1) *(coexistence).*
  b. *If γ/β ≥ ζ, z* = 0 *is the only convergence stable strategy (null provision).*
2. *If λ > ϱ, 𝒢*(*z*) *is unimodal in z with mode given by 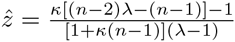. Furthermore:*
  a. *If γ/β ≤ ζ, both z* = 0 *and z* = 1 *are convergence unstable and there is a unique convergence stable strategy 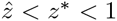(coexistence).*
  b. *If ζ < γ/β < η, there are two singular strategies z*_L_ *and z*_R_ *satisfying* 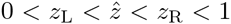. *The strategies z* = 0 *and z*_R_ *are convergence stable, whereas z*_L_ *and z* = 1 *are convergence unstable (bistable coexistence).*
  c. *If γ/β ≥ ζ, then z* = 0 *is the only convergence stable strategy (null provision).*

The arguments used for deriving this result are analogous to those used for deriving the results for club goods games with geometric production function (Result 2 in Appendix D). observing that *ϱ >* 1 holds for *κ ≥* 0 and that the case *λ* = 1 (constant returns to scale) is trivial, we can prove this result by considering three cases: (i) *λ <* 1, (ii) 1 *< λ ≤ ϱ*, and (iii) *ϱ < λ*.

For *λ <* 1, the production function is decelerating and hence Result 3 applies with Δ*β*_0_ = *β*. Recalling the definition of *ζ* = *κ*(*n* − 1) from (21) and rearranging, this yields Result 4.1 for the case *λ ≤* 1 *< ϱ*.

To obtain the result for the remaining two cases, we calculate the first and second forward differences of the benefit sequence (12) and substitute them into

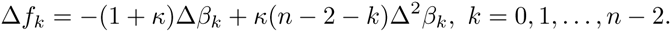

to obtain

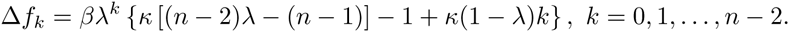

For *λ >* 1, the sequence Δ*f*_*k*_ is decreasing in *k* and hence can have at most one sign change. Moreover, since Δ*f*_*n-*2_ = *−βλ*^*n-*2^(1 + *κ*) *<* 0 always holds true, the sign pattern of Δ*f*_*k*_ depends exclusively on how Δ*f*_0_ = *β {κ* [(*n* − 2)*λ -* (*n* − 1)] − 1*}* compares to zero. observe, too, that *f*_*n*−1_ *<* 0 always holds true and that the sign of *f*_0_ is identical to the sign of *ζ − γ/β*.

Consider the case 1 *< λ ≤ ϱ*. Recalling the definition of ϱ (20) *we then have Δf*_0_ *≤* 0, implying that Δ*f*_*k*_ has no sign changes and that its initial sign is negative, i.e., *f*_*k*_ is nonincreasing. Hence, if *f*_0_ *≤* 0 (which holds if and only if *γ/β ≥ ζ*), the inclusive gain sequence has no sign changes and its initial sign is negative. otherwise, that is, if *γ/β < ζ* holds, we have *f*_0_ *>* 0 *> f*_*n*−1_ so that the inclusive gain sequence has one sign change and its initial sign is positive. Result 4.1 then follows from Peña et al. 2014, Result 3.

For *λ > ϱ* we have Δ*f*_0_ *>* 0, implying that Δ*f*_*k*_ has one sign change from + to −, i.e., *f*_*k*_ is unimodal. This implies that the gain function *𝒢*(*z*) is also unimodal with its mode 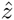 being determined by 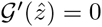 (Peña et al., 2014, Section 3.4.3). Using the assumption of geometric benefits, we can express *𝒢*(*z*) in closed form as

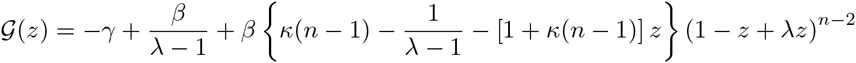

with corresponding derivative

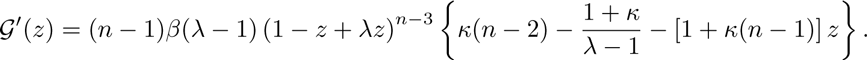

Solving 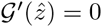 then yields 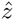 as given in Result 4.2. The corresponding maximal value of the gain function is

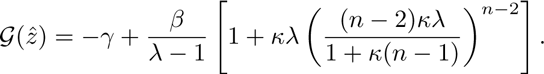

Result 4.2 follows from an application of Peña et al. 2014, Result 5 upon noticing that *f*_0_ *≥* 0 (precluding the stability of *z* = 0 and ensuring 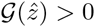 holds if and only if *γ/β ≤ ζ* and that 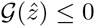 (ensuring that *B* dominates *A*) is satisfied if and only if *γ/β ≥ η*. (We note that the range of cost-to-benefit ratios *γ/β* for which bistable coexistence occurs is nonempty, that is *η > ζ* holds. otherwise there would exist a ratio *γ/β* satisfying both *γ/β ≤ ζ* and *γ/β ≥ η* which in light of Result 4.2.(a) and Result 4.2.(c) is impossible.)

## F other-only goods games

In other-only goods games, providers are automatically excluded from the consumption of the good they create, although they can still reap the benefits of goods created by other providers in their group. Payoffs for such games are given by *a*_*k*_ = *−γ* + *β*_*k*_ and *b*_*k*_ = *β*_*k*_, so that the inclusive gains from switching are given by *f*_*k*_ = *−γ* + *κ* [*k*Δ*β*_*k*−1_ + (*n* − 1 − *k*)Δ*β*_*k*_].

For this payoff constellation, it is straightforward to obtain the indirect benefits *𝔅*(*z*) from (B.3) in Appendix B. Indeed, observing that Δ*a*_*k*_ = Δ*b*_*k*_ = Δ*β*_*k*_ holds for all *k*, we have

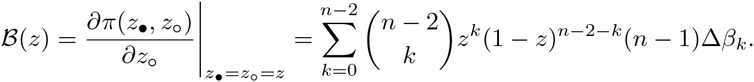

Using (9a) and the fact that *a*_*k*_ − *b*_*k*_ = *−γ*, we have that the direct benefit is given by −𝒞(*z*) = *−γ*.

Substituting these expressions for *𝒞*(*z*) and *𝔅*(*z*) into (4), we obtain

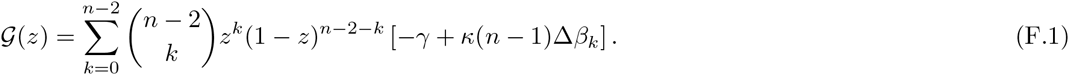

If *κ ≤* 0, our assumption that the production function *β*_*k*_ is increasing implies that *𝒢*(*z*) is always negative, so that *z* = 0 is the only convergence stable strategy (null provision).

To analyze the case where *κ ≥* 0, it is convenient to observe that (F.1) is of a similar form as (14). The only differences are that the summation in (F.1) extends from 0 to *n* − 2 (rather than to *n* − 1) and that the term multiplying the incremental benefit Δ*β*_*k*_ is given by *κ*(*n* − 1) (rather than by 1 + *κ*(*n* − 1)). All the results obtained for public goods games can thus be easily translated to the case of other-only goods games.

Specifically, we have the following characterization of the resulting evolutionary dynamics. With constant returns to scale, selection is frequency-independent with null provision if *κ < γ/*[(*n* − 1)*β*] and full provision if *κ > γ/*[(*n* − 1)*β*]. With diminishing returns to scale, the gain function is decreasing in *z* (negative frequency dependence). There is null provision if *γ ≥ κ*(*n* − 1)Δ*β*_0_, and full provision if *γ ≤ κ*(*n* − 1)Δ*β*_*n-*2_. If *κ*(*n* − 1)Δ*β*_*n-*2_ *< γ < κ*(*n* − 1)Δ*β*_0_ holds, there is coexistence. With increasing returns to scale, the gain function is increasing in *z* (positive frequency dependence). There is null provision if *γ ≥ κ*(*n* − 1)Δ*β*_*n-*2_, and full provision if *γ ≤ κ*(*n* − 1)Δ*β*_0_. If *κ*(*n* − 1)Δ*β*_0_ *< γ < κ*(*n* − 1)Δ*β*_*n-*2_, there is bistability.

If the production function is geometric (12), the gain function is given by

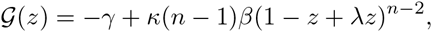

so that, for *λ /*= 1, the evolutionary dynamics are similar to the case of public goods games after redefining the critical cost-to-benefit ratios as

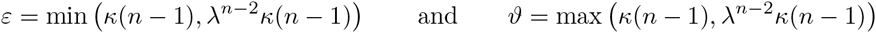

and letting

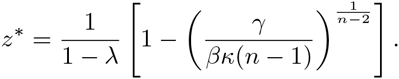

